# PySteMoDA: An Open-Source Python Package for the Analysis of Steered Molecular Dynamics Simulations Data

**DOI:** 10.64898/2026.01.26.699872

**Authors:** Ismahene Mesbah, Colin Klaus, Marcos Sotomayor, Fidan Sumbul, Felix Rico

## Abstract

Molecular dynamics simulation is a powerful computational technique used for predicting and understanding the dynamic behavior of biomolecular systems. Steered molecular dynamics (SMD) simulations enable the study of force-induced processes in biomolecules, effectively mimicking single-molecule force spectroscopy experiments probing protein unfolding and receptor-ligand unbinding. Given the stochastic nature of these mechanical events, accurately exploring the dynamic behavior of biomolecules and extracting accurate physical information requires several *in-silico* experiments. This includes performing many pulling simulations at different velocities or force loading rates. The large amount of data obtained from these simulation sets requires efficient automated data processing tools. We present PySteMoDA, a novel Python package with a user-friendly graphical interface specifically designed for constant-velocity SMD data analysis. The automated force peak detection methods reduce user bias, improve accuracy, and accelerate data analysis. The package also allows identification of residues involved in mechanical events through computation of the time-dependent mechanical work and correlation factors between residue pairs. This package not only addresses automated data processing in SMD simulations and accurate parameter extraction, but also significantly enhances accessibility and usability. Through PySteMoDA, users can efficiently analyze simulation data without the barrier of coding, facilitating a wider range of investigations and insights in the field of computational biochemistry and biophysics.

## Introduction

Molecular dynamics (MD) simulations used as “computational microscopes” enable atomic-level prediction and understanding of biomolecular processes such as side chain fluctuations and conformational dynamics of proteins (1–6). MD simulations of biomolecules are commonly carried out using packages such as CHARMM (7), OpenMM (8), AMBER (7), GROMACS (8), and NAMD (9). NAMD is open source, easy to use, highly optimized for massively parallel supercomputers, and compatible with the visual molecular dynamics (VMD) software (10) that permits interactive analysis of simulation outputs. Steered molecular dynamics (SMD) simulations represent a powerful computational technique that mimics single-molecule force spectroscopy (SMFS) experiments and, thus, allows for the exploration of molecular interactions and conformational changes during relevant biological processes where mechanical force is involved (11–14). By applying external forces to specific atoms or molecules, SMD enables the study of processes such as protein unfolding, ligand-receptor unbinding, and the mechanical probing of macromolecular structures (15,16). In addition, SMD can be used to accelerate processes that otherwise occur over timescales beyond those accessible to standard equilibrium MD simulations (17). The application of SMD has been instrumental in elucidating the pathways and energetics of mechanically induced conformational transitions and intermediate states that are often critical to the function of biomolecules. For example, simulations of mechanical unfolding in proteins—such as titin in muscle sarcomeres (16)—and ligand-receptor unbinding events, like (strept)avidin and biotin (15,18,19), offer valuable insights into the mechanisms and forces underlying these processes, including those that involve highly stable interactions (20). The ability to model these forces and to predict and understand their effects at a microscopic level is essential for the design of new drugs, the engineering of protein-based materials, and the development of nanoscale devices.

SMFS experiments probe the mechanical properties of biomolecules, while SMD simulations provide atomistic insight into their force-induced responses. For instance, SMD revealed that unfolding of the titin I27 (I91) domain begins with separation of the A and B β-strands, followed by disruption of the A’–G strands (16,21). Early SMD studies were limited to pico-to nanosecond timescales, which made direct comparison with slower SMFS experiments difficult (18,22). Despite this, their molecular predictions were later validated at millisecond timescales (23). Advances in high-speed SMFS and long-timescale SMD now allow overlapping timescales (19,24), and many relevant mechanical events occur within microseconds (25–29). This timescale is accessible with modern MD engines and supercomputers (2,30).

In SMD simulations, the pulling velocity or force is kept constant. In constant-velocity simulations a virtual spring is attached to an atom or group of atoms, and the free end of this spring is moved at a constant velocity, applying a force proportional to the spring extension that induces conformational changes in the biomolecule. In constant-force SMD simulations a force is directly applied to an atom or group of atoms in a predetermined direction. In both constant-velocity and constant-force simulations, the protein is either stretched from one end by keeping the other end fixed or from both ends in opposite directions. Other configurations for force application, including length clamp (31), can be implemented as well.

As a typical constant-velocity SMD simulation progresses, the force applied to the molecule and atomic coordinates of each atom are recorded at defined time intervals. Different parameters are then extracted from simulation trajectories, including unfolding forces, molecular extensions, molecular stiffnesses and loading rates. For many systems, it is now possible with modern computers to carry out hundreds of simulations for more accurate statistical analysis of these dynamic ensembles (19,24,32–34). However, manually analyzing the large amount of data generated by SMD simulations is time-consuming and prone to error. To address this challenge, we introduce PySteMoDA, a Python package designed to semi-automatically extract and analyze data from SMD simulations.

PySteMoDA is an open-source, freely accessible, semi-automated, and user-friendly package that facilitates the analysis of the output files of constant-velocity SMD simulations performed using NAMD. It features a graphical user interface (GUI) allowing users to carry out advanced analysis without needing to write code. Additionally, a Google Colab version is available, enabling users to run the pipeline directly without local installation. Once configured, the entire pipeline, including data processing, peak detection, loading rate calculation, correlation analysis, and inter-residues deformation measurement, runs automatically using the default values provided by the platform. Both the GUI- and Colab-based implementations significantly lower the barrier to entry and improve usability for researchers across disciplines interested in SMD analysis.

## Material and Methods

### 1. Implementation

PySteMoDA was developed in Python 3.8 and requires several package dependencies including Tkinter (35), ProDy (36), scikit-learn (37) and SciPy (38). Our package adopts an object-oriented model, enhancing its scalability, modularity, and reusability. This design choice enables users to efficiently extend its functionality to suit their specific research needs. A user-friendly GUI ensures that even newcomers to the field can quickly become proficient with PySteMoDA. This simplicity enables users to save time and easily perform their analysis. The package is executable on Linux, Mac, and Windows operating systems. The PySteMoDA workflow involves inputting file names, extracting data, doing analyses and exporting results from analyses (**Figure 1)**. Various Jupyter notebooks have also been implemented to allow batch processing from the command line. Additionally, a Google Colab version is available, enabling users to run the pipeline directly without local installation.

**Figure 1:**
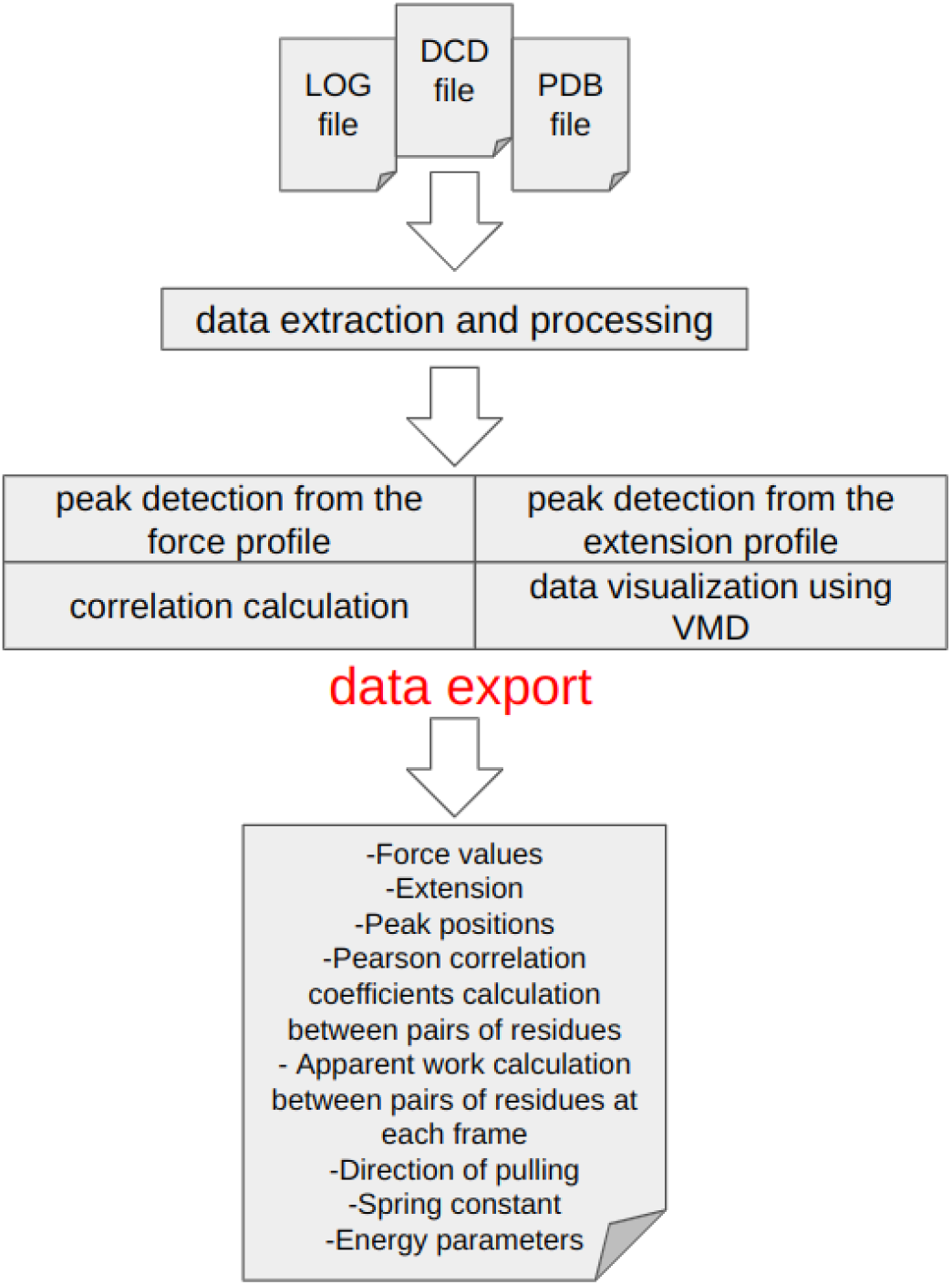
Schematic representation showing the PySteMoDA workflow. Analysis starts by loading the LOG and DCD files obtained from SMD simulations using NAMD in addition to the PDB file containing the atomic coordinates. Loading a protein structure file (PSF) is recommended to keep initial connectivity. Users can perform peak detection either from the force or extension curves. The Pearson coefficient of correlation and apparent work is calculated between pairs of residues at different frames, which allows users to observe the effect of changes in distance between residues on force. The simulation movie can be visualized via VMD where the DCD and PDB files are uploaded automatically by PySteMoDA. At the end of the analysis, users have the possibility to export the analysed data in different formats.

### 2. Simulations

Representative data used to validate and showcase PySteMoDA in this manuscript were obtained from two sets of NAMD SMD simulations. The first set corresponds to SMD simulations of a cadherin-23 (CDH23) subdomain including extracellular cadherin (EC) repeats EC19-20 with three bound calcium ions (extracted from PDB: 5TFK and hereafter referred to as CDH23 EC19-20) (39,40). Details of these simulations are described elsewhere. Briefly, simulations of the CDH23 EC19-20 system (*n*_*atoms*_ = 244,076 atoms) were performed using NAMD 2.12 (9) with the CHARMM36 force field (41) and the TIP3P water model. Van der Waals forces were computed using a 12 Å cutoff with a switching function starting at 10 Å. Electrostatic forces were computed using the particle mesh Ewald algorithm with a grid spacing of 1 Å. Dynamics in time was advanced using a 2 fs timestep. The system was first minimized for 1,000 steps, then subjected to 50,000 steps of constrained backbone dynamics (*k* ∼ 700 pN/nm). A Langevin thermostat and hybrid Nosé-Hoover piston method were used to maintain a temperature of 300 K and pressure of 1 atmosphere (*NpT* ensemble). In the first 1 ns a Langevin damping coefficient of 1 ps^-1^ was used for the thermostat while in the following 10 ns a Langevin damping coefficient of 0.1 ps^-1^ was used. A piston period of 200 fs and a damping timescale of 100 fs were used for pressure control. For SMD simulations, virtual springs were attached to the N- and C-terminal C_α_ atoms with spring constants of *k* ∼ 700 pN/nm. The free ends of the virtual springs were moved at stretching speeds of 5 nm/ns in opposite directions (10 nm/ns total). SMD data output and coordinates were written to disk every 40 fs and every 1 ps, respectively.

The second set of simulations used to validate and showcase PySteMoDA involved the intercellular adhesion molecule 1 (ICAM-1, *n*_*atoms*_ = 67,215) which was simulated with NAMD 2.9 (9) using the CHARMM36 force field (41,42) and the TIP3P water model. Dynamics in time were advanced using a 2 fs timestep. The results are reported in the supplementary material. We modelled ICAM-1 based on the cryo-EM structure of D1-D5 (PDB ID: 1Z7Z) (43) and crystal structure of D3-D5 (PDB ID: 2OZ4) (44). Disulfide bonds were specified using the corresponding patch to create the protein structure file (PSF) for simulation. ICAM-1 was stretched at 1 nm/ns along a direction defined by the vector that connected the fixed and SMD atoms after equilibration (*v* = 0. 0524, 0. 3795, 0. 9236). The N-terminal Cα atom was fixed during the simulation. The spring constant corresponds to 104 pN/nm. The simulation was carried out over a total of 70 ns with outputs written every 10 ps for both DCD (trajectory coordinates) and SMD frequencies. In the case of the ICAM-1, the simulation was conducted in the *NVE* ensemble, where the number of particles, volume, and total energy were conserved throughout the run.

### 3. Data extraction and conversion

PySteMoDA requires NAMD output files, including the standard output (generally saved to a file with extension LOG) and a trajectory DCD file. These files are used to extract data, including external forces applied during the simulation to manipulate the molecule, commonly referred to as “SMD forces” **(See supp. information)**. These are the forces used to mimic either physiological or experimental conditions such as those used in atomic force microscopy (AFM) and optical tweezers (OT). External forces can be applied to atoms in NAMD simulations in different ways, including by using the SMD module, the “tool command language” TCL interface, or the collective variables module. PySteMoDA can read the NAMD LOG file assuming the standard SMD output format (“SMD” *timestep x y z f*_*x*_ *f*_*y*_ *f*_*z*_) used for simulations in which one set of atoms is stretched. For simulations in which two groups of atoms are stretched, PySteMoDA assumes that one group is stretched using the SMD module and the other is stretched using the TCL interface with a predefined output format (“TCL:” *timestep x y z f*_*x*_). Both SMD and TCL forces are extracted **(See supp. information)** and can be visualized within the GUI with respect to simulation time or stretching distance. The stretching distance corresponds to the movement of the SMD atom.

PySteMoDA also extracts atom positions and time steps from the trajectory DCD file. The package calculates the extensions at each frame that the user provides in the trajectory file. These correspond to the Euclidean distances between the Cα atoms of two given residues *r*1 and *r*2 based on their three-dimensional coordinates at each time step, according to:

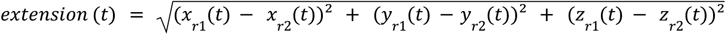

The extension is calculated by default at all the frames between the Cα of the first residue (*r1*) and the Cα of the last residue (*r2*), although users can select any other residue or frame coordinates.

The correlation matrix of forces and distances between pairs of residues is obtained by calculating the Pearson coefficient of correlation. To achieve this, first, PySteMoDA calculates the center of mass (COM) of each amino acid residue for each Cartesian coordinate as:

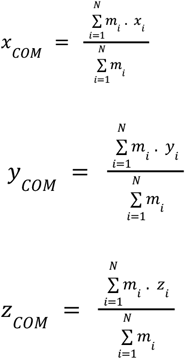

where *x*_*i*_, *y*_*i*_, *z*_*i*_ are the cartesian coordinates of the ith atom, *m*_*i*_ is its atomic mass, and *N* is the number of atoms. PySteMoDA uses the COM coordinates (instead of Cα coordinates) because these represent the average position of all the atoms in a residue, weighted by their masses. This provides a single location that reflects the distribution of the entire molecular mass. Therefore, using the center of mass gives a more global, holistic view of the residue’s position and movement while Cα coordinates represent only the backbone of the protein. Once the centers of mass of residues are obtained at each frame (*k*), their Euclidean distances are computed and used as the distance between pairs of residues (*r*1, *r*2):

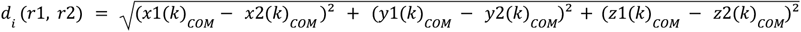

where *x1, y1, z1* are the spatial coordinates of *r1* and *x2, y2, z2* are the coordinate of *r2*. The Pearson correlation “work” coefficient is calculated between forces and distances for each pair of residues that are within 13 Å (default value that can be manually changed) at a given range of *n* frames from:

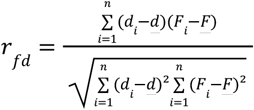

where *d*_*i*_ is the distance between the pair of residues at frame *k, d* is the mean distance between this pair of residues along the range of frames [*k:k+n*], ***F***_*i*_ is the force vector at frame *k*, and *F* is the mean force along the given range of frames [*k :k+n*].

The **apparent work** (in Joules) is calculated along the trajectory frames via the dot product of the obtained distances vector (Δ*r*) and the force vector (direction of pulling):

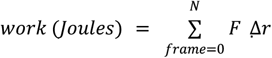

where ***F*** and *Δ****r*** are the force and distance vectors. The Δ*r* vector is calculated for each pair of residues using the center of mass:

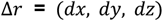

where :

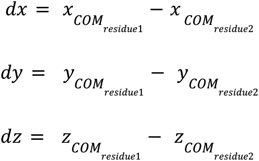

### 4. Detection and characterization of unfolding/unbinding events

Detection of unfolding/unbinding events was implemented using two different approaches:

#### 4.1 Smoothing and automatic peak detection using the force curve

In this approach, an algorithm scans the force curve to identify local maxima. A local maximum is a point where the value is greater than its neighboring points. These local maxima represent potential specific unfolding/unbinding events in the data. For each identified local maximum, the prominence is calculated where prominence is defined by NumPy’s find_peaks function. Prominence is a measure of how much a peak stands out from the neighboring peaks. Peaks with a prominence below a certain threshold are considered as noise and are filtered out by PySteMoDA. By default, this threshold is computed automatically as the mean value of all peak prominences. This default thresholding step helps eliminate smaller peaks that may not be of interest. Users can modify this threshold value if a different level of sensitivity is desired.

The interface offers a “natural cubic spline” smoothing filter. In this context, knots are specific points along the *x*-axis where individual cubic polynomial pieces meet and are joined together. A natural cubic spline fits the data by dividing the curve into intervals (or windows) defined by these knots. Within each interval, a cubic polynomial is used, and at each knot the spline is constructed so that the function, its first derivative, and its second derivative are all continuous. The number of knots can be defined by the user or automatically filled within the interface (with knots spaced at regular intervals of approximately 0.3 nm along the *x*-axis). The number of knots directly controls the flexibility of the spline, for example, more knots allow the spline to follow fine-scale variations in the data, while fewer knots result in a smoother, more global fit.

#### 4.2 Semi-automatic peak detection using the extension curve

In this approach, the derivative of the extension-time curve is calculated and plotted. This mathematical operation involves determining the rate of change of extension with respect to time, and the resulting derivative graph serves to detect sharp transitions in the original curve as the increasing steps in the extension appear as sharp peaks in its derivative. The process of calculating the derivative facilitates a more precise and efficient detection of their locations.

#### 4.3 Peak analysis

Once the peaks are detected, relevant parameters such as the loading rate (pN/s) and molecular and effective stiffness are computed automatically, within a given extension window before the peak (1 nm by default, but manually modifiable). The loading rate is determined from the slope of the force versus time signal over the 1 nm default distance before the peak and the effective stiffness as the loading rate divided by the pulling velocity. The molecular stiffness is calculated using the formula:

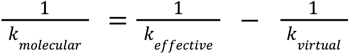

where *k*_*molecular*_ is the molecular stiffness, *k* is the effective stiffness, and *k* _*effective virtual*_ is the virtual spring constant.

### 5. Inter-residues distance deformation

Finally, an inter-residues distance deformation is calculated. This metric is a useful way to track structural changes in the protein over time at the residue level. It, therefore, provides a measure of how much the spatial arrangement of amino acids deviates from its original configuration.

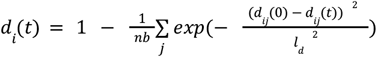

The formula computes a deformation metric that quantifies the change in the distances between residues in a protein over time. Here, *d*_*ij*_ (0) is the initial distance between residue *I* and *j* at frame 0, *d*_*ij*_ (*t*) is the distance between them at a later time *t*, and *l*_*d*_ is a characteristic length scale (usual distance between two amino acids) used here to normalize the obtained value. The sum runs over all residue pairs *i,j*, and *nb* represents the total number of pairs being considered. If the distance between amino acids is large the term 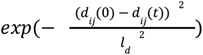 results in smaller values (i.e., more deformation). The final result reflects how much the distances between residues have changed on average over the specified frames.

This formula was adapted to incorporate mechanical work and, thus, to analyze the work due to applied forces:

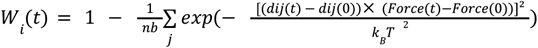

where:

*Force*: is the applied force in pN.

*k*_*B*_*T*: is the Boltzmann constant multiplied by the temperature.

The distance deformation measurements can be carried out within the google colab but can also be calculated using the command line. After opening a terminal, users can use the following command to obtain the deformation measurement:

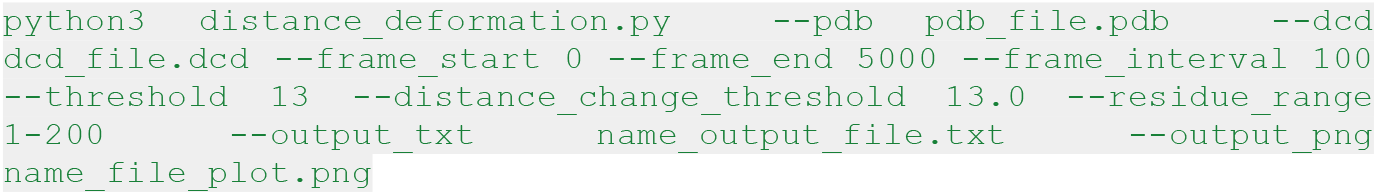

The “threshold” parameter here is the maximum distance to be considered when calculating the deformation metric, the “distance change threshold” parameter is the maximum distance to report in the “output_file” once the residues separate from each other during the pulling.

## Results

PySteMoDA is freely available and downloadable (https://gitlab.com/fm4b_lab/pystemoda) and can also be accessed as a Google Colab notebook (https://colab.research.google.com/drive/1Cc1IqlMnsJ-SDoE3CIiMhnLHVN5y61Dp?usp=sharing). To illustrate PySteMoDA’s user-friendly GUI we describe here various steps to carry out analyses of SMD simulations for two systems, CDH23 EC19-20 and ICAM-1. We will focus first on standard steps for the analysis of CDH23 EC19-20 SMD simulations and will finish with a deformation analysis for ICAM-1.

The first step consists of uploading the NAMD output (LOG) file of the simulation. PySteMoDA extracts relevant data from it and converts units automatically. The force versus displacement (also called distance) for CDH23 EC19-20 simulations shows one clear main peak followed by a minor peak at both ends (SMD and TCL) of the protein fragment (**Figure 2**). Users can also plot and analyze different quantities such as temperature, pressure, and VDW energies. Other data can be exported, including force (pN), virtual spring constant (pN/nm), velocity (nm/ns), displacement/distance (nm) and time (s). PySteMoDA allows plotting and analysing both SMD and TCL forces when available by simply checking the “*See F(-v)*” button (**Figure 2**). Moreover, PySteMoDA allows the users to smooth the thermal noise by applying a natural cubic spline smoothing filter (45) that requires specification of a “knots” parameter. PySteMoDA provides a default value of “knots” to guide users (**see Methods**). Smoothing of the CDH23 EC19-20 SMD data after applying the natural cubic spline filter with the suggested default number of “knots” shows clearer force peaks (**Figure 3A**). These automated data extraction tools are a key benefit of PySteMoDA.

**Figure 2:**
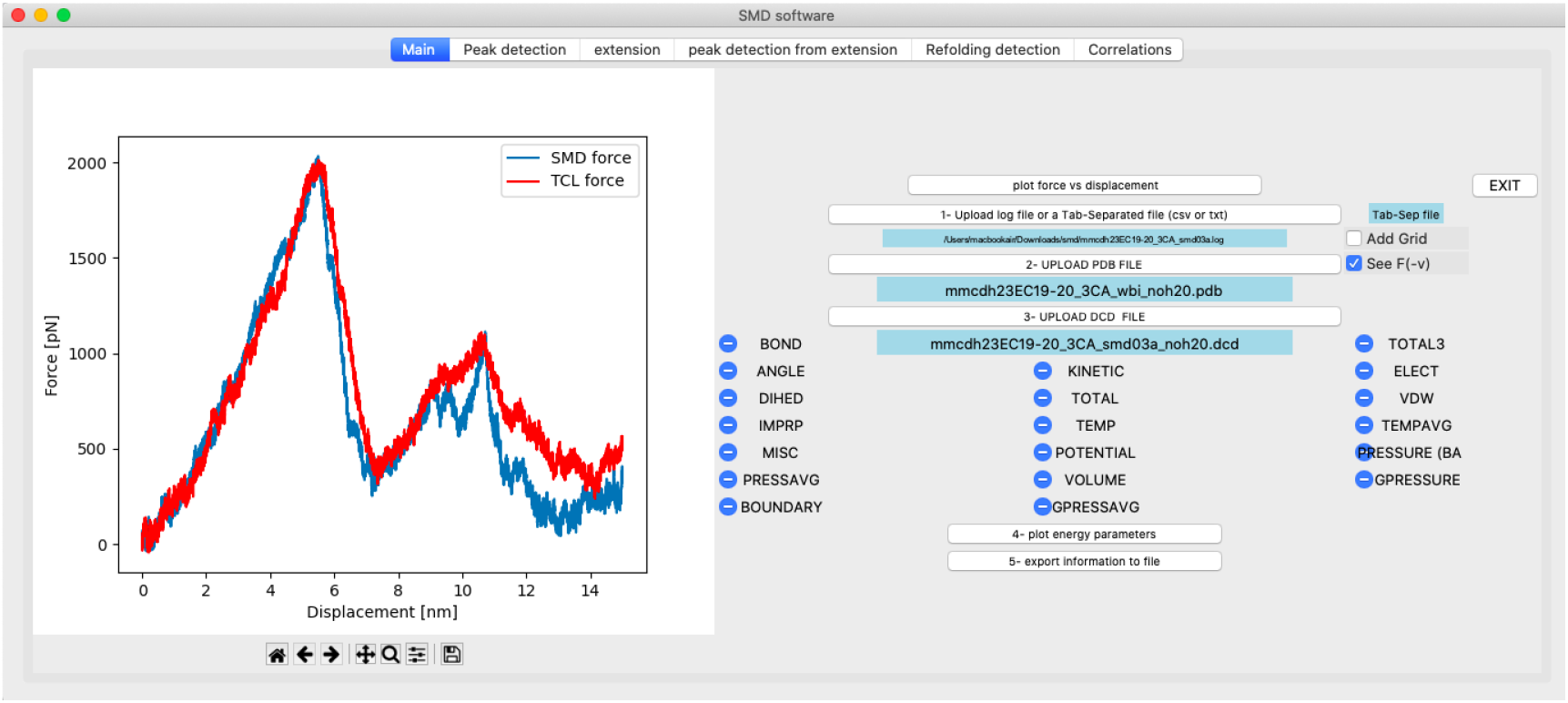
Snapshot of the PySteMoDA main tab. The force-displacement (also called force-distance) curve for the extracellular cadherin (EC) fragments 19 and 20 of cadherin-23 with three calcium ions (CDH23 EC19-20) is shown for a SMD simulation at 10 nm/ns. In this example, the protein was pulled from both ends, the TCL force curve is shown in red and the SMD force in blue. TCL forces can be visualized by checking the “See F(-v)” button.

**Figure 3:**
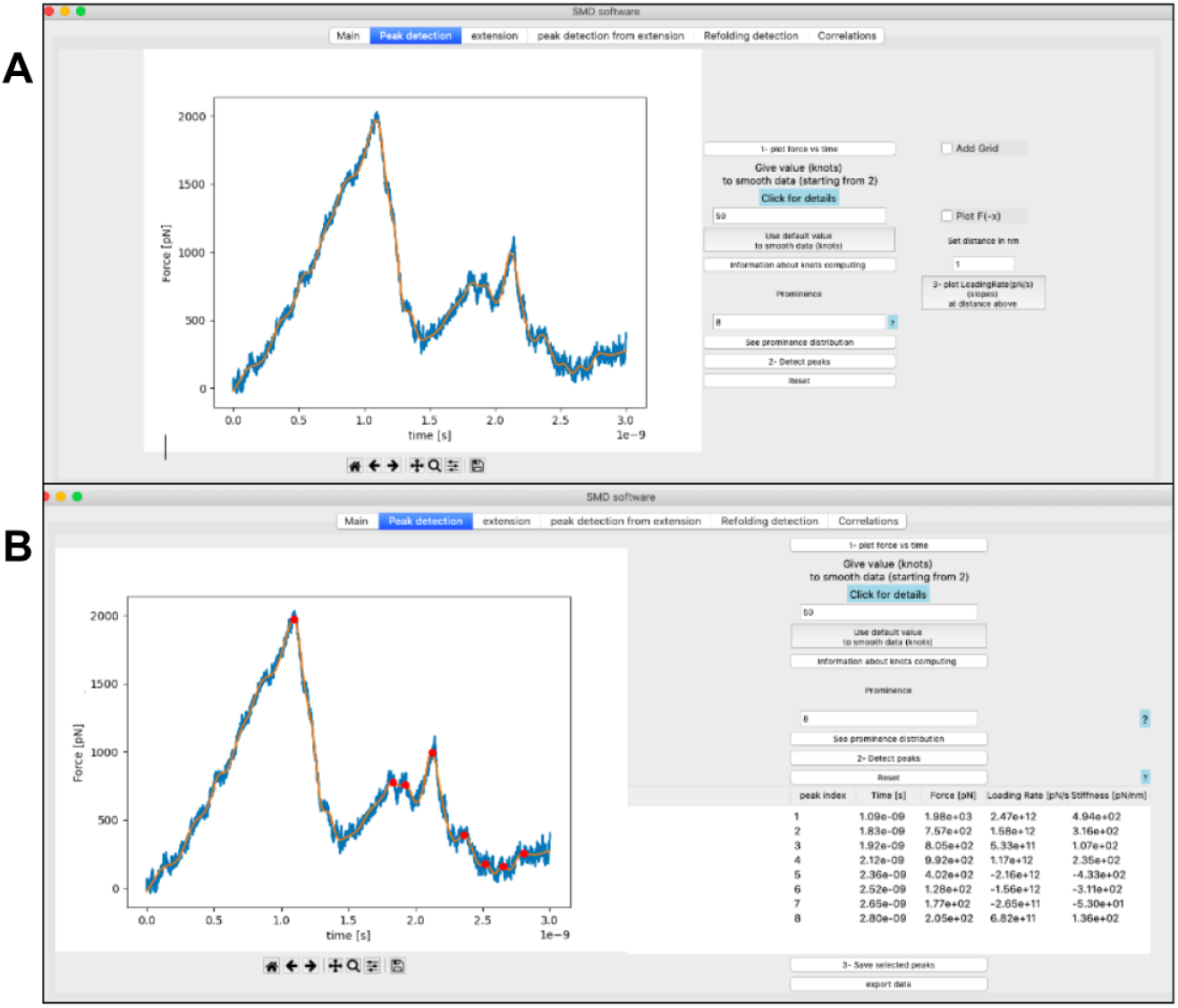
Data smoothing and peak detection. **(A)** Snapshot showing the natural cubic spline smoothing filter applied with default values to the force curve(orange, filtered; blue, raw). **(B)** Snapshot of the tab dedicated to peak detection from the force curveof the unfolding of CDH23 EC19-20 using the default parameters for smoothing (knots) and prominence. The list of the corresponding peaks is displayed within the table where the time (s), force (pN), loading rate (pN/s) and stiffness (pN/nm) associated to each peak are automatically calculated and displayed.

In addition, PySteMoDA facilitates the analysis of SMD trajectories by enabling detection of specific unfolding/unbinding peaks from the force curve. To optimize this process, default values of relevant parameters, such as the prominence threshold, are suggested and automatically filled within the interface as soon as data are uploaded. Peak detection using the prominence parameter helps to identify sharp peaks that should correspond to unfolding/unbinding events. Peaks with high prominence are considered distinct from the background noise. The prominence value suggested by PySteMoDA corresponds to the mean prominence of all the potential peaks in the curve (**See Methods**). Users can modify the default values of knots and prominence if not satisfied with the output. The peak detection algorithm identifies 8 unfolding events for the stretching of the CDH23 EC19-20 protein fragment using the default values of smoothing (knots) as well as the prominence value (**Figure 3B**). All the detected peaks are displayed in a table that records unfolding/unbinding forces (pN), time (s), loading rates (pN/s), stiffnesses (pN/nm) and molecular extensions (nm) **(Figure 3B)**. Users can select the relevant peaks and export the information displayed within the table to different file formats (.csv, .dat, or .txt) **(supp. Table S1**). We expect these PySteMoDA features to facilitate analyses of SMD trajectories.

Calculation of the loading rate at the main unfolding peak of the CDH23 EC19-20 molecule reveals a value of 2.5 x 10^12^ pN/s, as expected for a simulation done at a very fast stretching speed (10 nm/ns) using a stiff spring constant (**Figure 4**). The loading rate is obtained via a linear fit of the force-time curve and plotted at each peak within the GUI. The linear fit is calculated at a distance of 1 nm before each peak. This default distance was shown to work well on the example simulations but can be modified within the corresponding entry **(Figure 4)**. Peak detection and corresponding analyses can also be done for TCL forces when these are applied during the SMD simulation **(supp. Figure S2)**.

**Figure 4:**
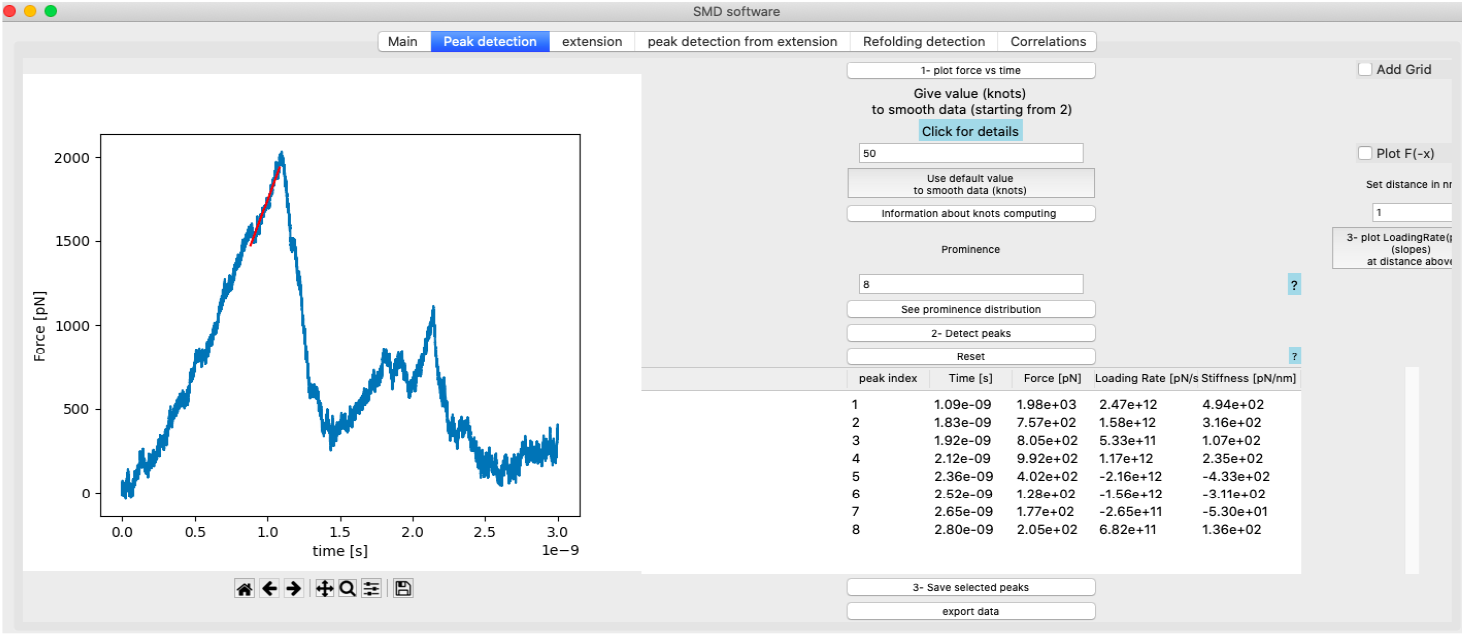
Loading rate calculation. Snapshot of loading rate (pN/s) plot displayed within the GUI after calculating the slope of the linear regression fit at the peak in the force versus time curve of CDH23 EC19-20. This plot appears when the user clicks the “plot loading rate” button while the loading rate (pN/s) and stiffness (pN/nm) values are calculated automatically and displayed within the main table shown within the GUI.

PySteMoDA can extract the extension curve corresponding to the Euclidean distance between the N- and C-terminal residues at each frame, which corresponds to the stretching of the molecule. When plotting the extension-time curve the unfolding events may appear as deviations from a linear increase in extension (**Fig. 5A**). The extension curve of CDH23 EC19-20 in nm with respect to time in seconds shows two unfolding events as deviations from a linear increase in extension that resemble steps (**Figure 5A**). When these steps are sharp, they can be accurately detected from the derivative of the extension curve. This alternative peak detection approach from the time derivative of the extension is provided by PySteMoDA and available within the “*peak detection from extension*” tab. Within this tab, users can identify the unfolding events by selecting the number of highest peaks *N*. PySteMoDA identifies two unfolding events in the derivative of the extension profile of CDH23 EC19-20 (**Figure 5B**). These peaks can be selected by the user and represented in the force vs time curve (**Figure 5C**). Extension curves can also be calculated across a given range or ranges of amino acids (separated by commas, e.g. 1-89, 90-165…), for example, to define the extension of each domain in a multidomain protein (**Figure S3**). We expect this extraction pipeline to accelerate analyses of similar SMD trajectories and to easily correlate force peaks with protein regions undergoing unfolding.

**Figure 5:**
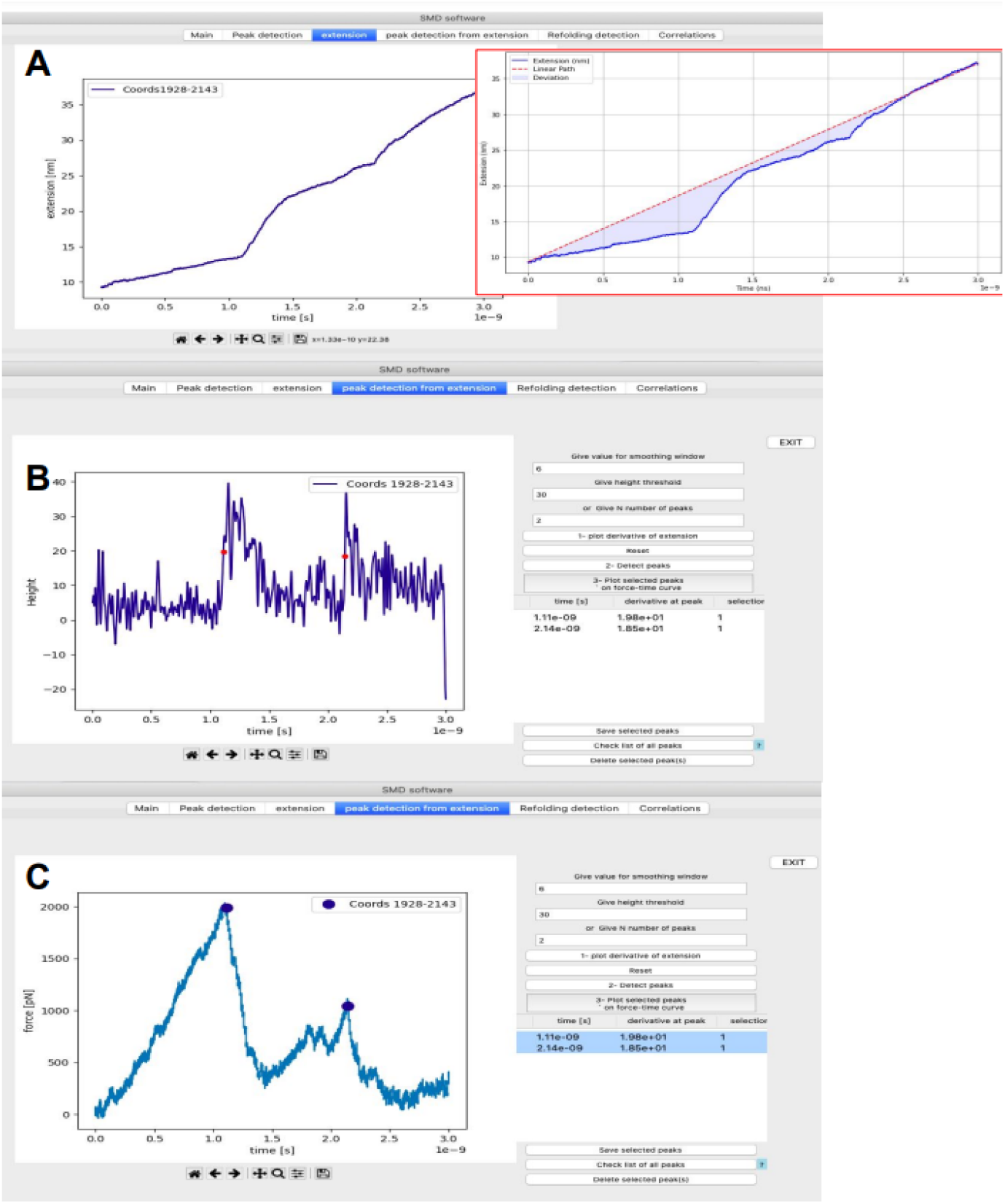
Snapshot of the *peak detection from extension* tab for CDH23 EC19-20. **(A)** Extension vs time curve. The unfolding peaks appear as increasing steps in extension. Inset shows linear change of position for free end of the spring. **(B)** Derivative of the extension facilitates peak detection. A Gaussian filter has been applied to smooth the derivative of the data. The input parameters to detect the unfolding peaks are height and the number of highest peaks (*N)*, which were set to 30 and 2 respectively. **(C)** The two peaks detected in **(B)** are represented by circles in the force versus time curve.

PySteMoDA can calculate a matrix of correlations to gain insight into how distances between residue pairs could determine changes in force during unfolding/unbinding processes. For this, the Pearson coefficient is calculated between the distances of each residue pair and the forces. This correlation matrix is represented as a heatmap within the graphical interface. The correlation heatmap for the CDH23 EC19-20 protein fragment shows positive correlation coefficients in red and negative coefficients in blue (**Figure 6**). The heatmap can be exported as a static image (png, jpeg and svg) or a dynamic movie that shows the correlations every *N* trajectory step defined by the user. The file format in this case is either .gif or .mp4, both available as outputs within the GUI and the Colab notebook. The correlation coefficient values can also be exported as a matrix within a file (.txt, .csv, .dat). These coefficients of correlation are calculated along a range of time frames. By default, PySteMoDA uses the range from the first to last time frame, automatically extracted from the DCD trajectory file. This default range can be changed as needed, e.g., to select the frames around the major unfolding event. The residues taken by default in the correlation calculation are the first and last residue, their corresponding coordinates (*x, y, z*) are automatically extracted by PySteMoDA from the PDB file. The work (Joules) heatmap is also calculated along the indicated range of frames. An example of the work heatmap before and after the major unfolding peak shows that CDH23 EC20 has partially unfolded (**Figure 7**). The residue range can also be modified and can be provided in broken ranges, which may be useful for large protein structures with multiple domains. These analyses can be used to provide insight into which residues are most correlated with force rupture events.

**Figure 6:**
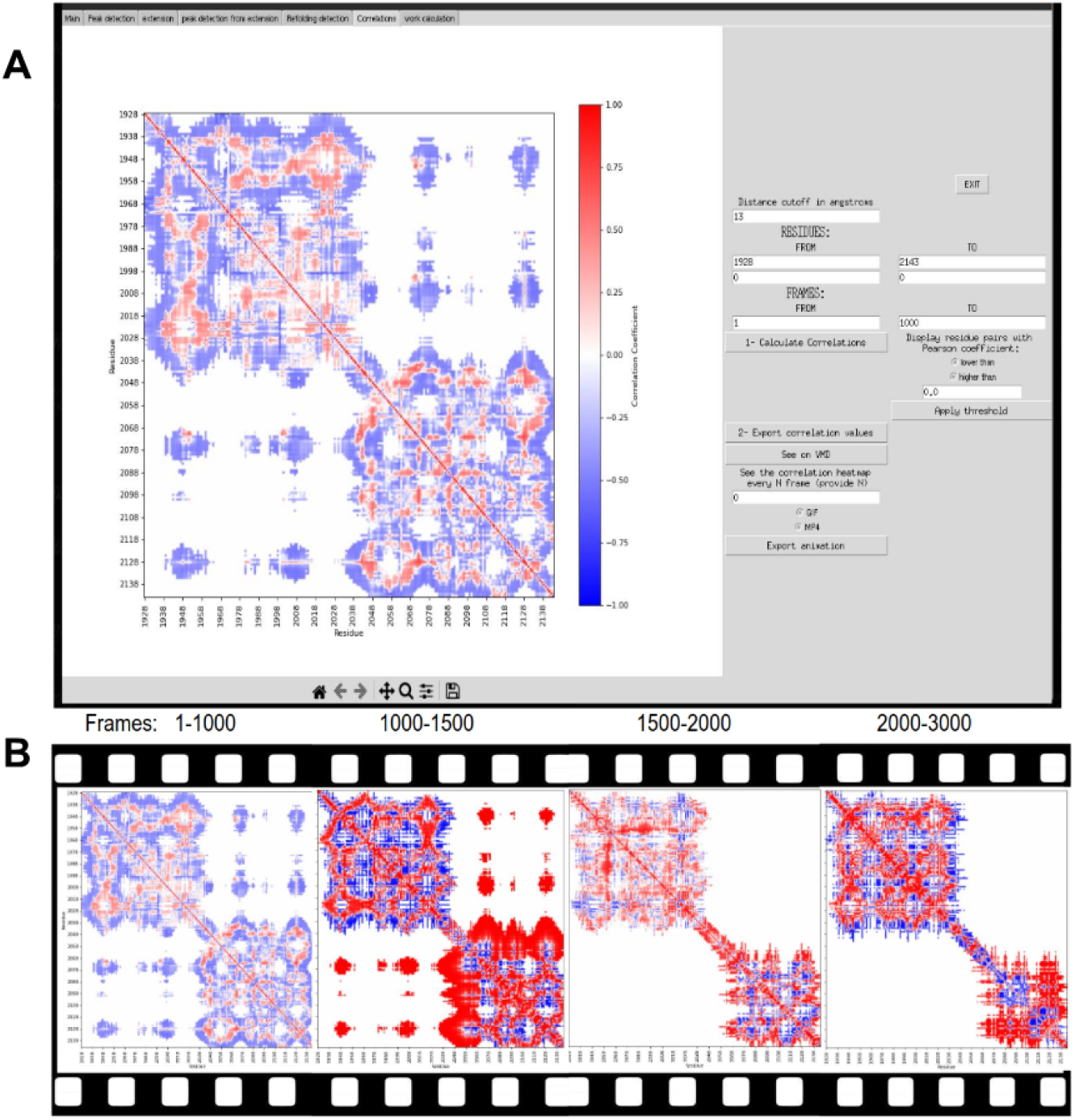
Pearson correlation computation. **(A)** Snapshot of the heatmap that shows the Pearson coefficients of correlations between force and all pairs of residues of CDH23 EC19-20 within a distance threshold of 13 Å (can be modified by the user) between the frames 1-1000. The correlation matrix within the interface is shown with the corresponding parameters. Note that the coefficient is equal to 0 when the color is white, negative values are blue, and positive values are red. **(B)** The correlation map along the unfolding trajectory. The first unfolding peak occurs after frame 1,000 (i.e., after 1 ns) where we see positive correlations (red) in EC20. This indicates that when the force is increasing the distance between pairs of residues is also increasing. In other words, the EC20 fragment is unfolding.

**Figure 7:**
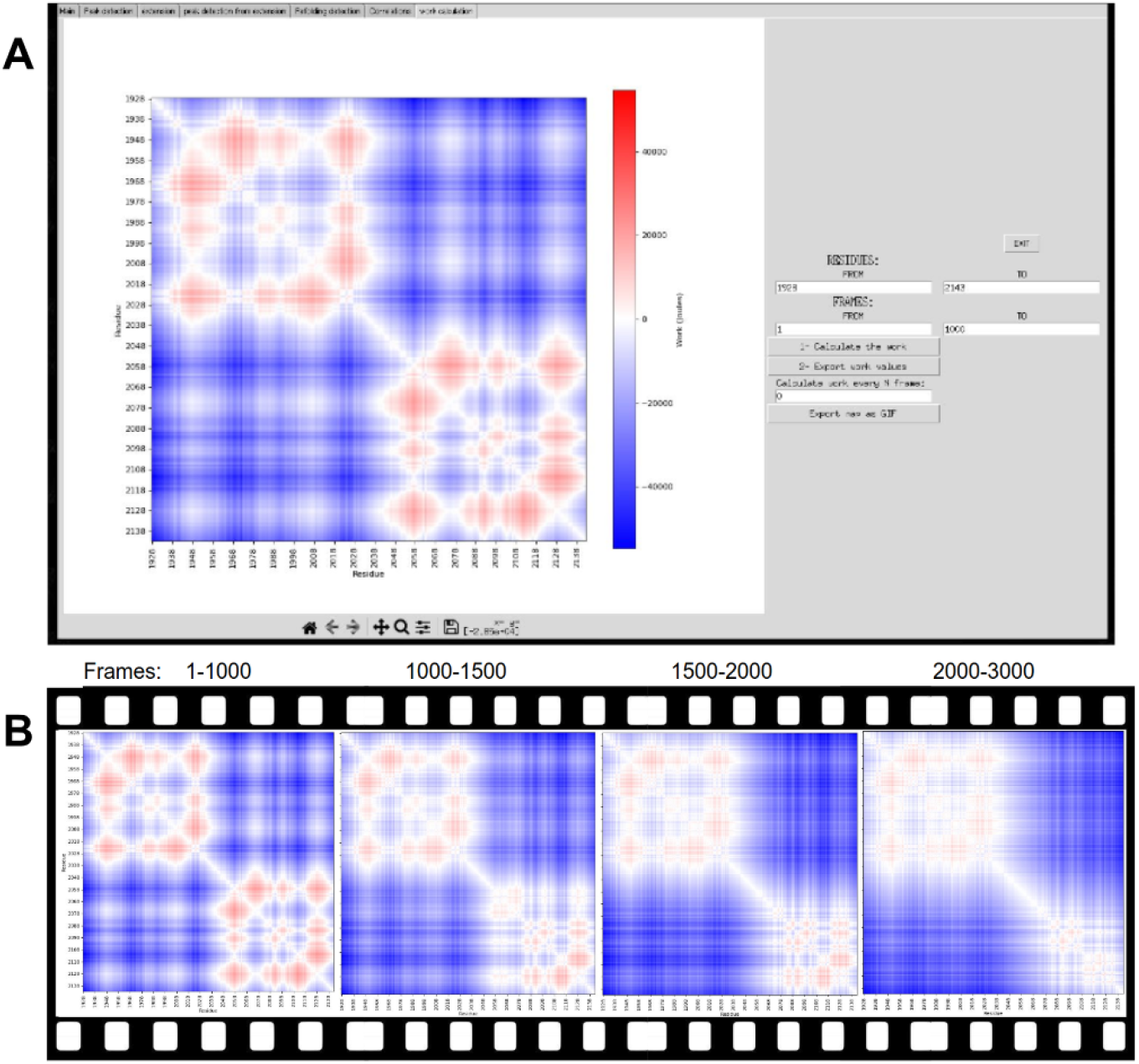
Heatmap of work for each pair of residues along the unfolding trajectory of CDH23 EC19-20. Negative work values in Joules appear in blue, positive values in red while the white color corresponds to zero. **(A)** snapshots of the heatmap of work within the interface with the corresponding parameters (i.e, frame ranges). **(B)** the heatmap of the work between frame 1 and 3,000 (i.e., from 0 to 3 ns). It is visible in the last map that EC20 has partially unfolded although the C-terminal end of the fragment is still resistant to force.

PySteMoDA can also be used to compute the inter-residue distance deformation, which was done here to characterize the mechanical response of the fourth domain of ICAM-1 (residues 282–364) during stretching in SMD simulations (**Figure 8**). The distance deformation matrix was calculated every 50 simulation frames, with frame intervals reported along the *y*-axis and residue numbers along the *x*-axis. Distinct clusters of residues undergoing unfolding events are observed as regions highlighted in light green and yellow at specific frame intervals. The temporal evolution of the extension of the fourth ICAM-1 domain (red) and the total applied force (blue) shows correlated changes over the same time scale. This enables direct association between residue-level unfolding events, domain extension, and force response.

**Figure 8:**
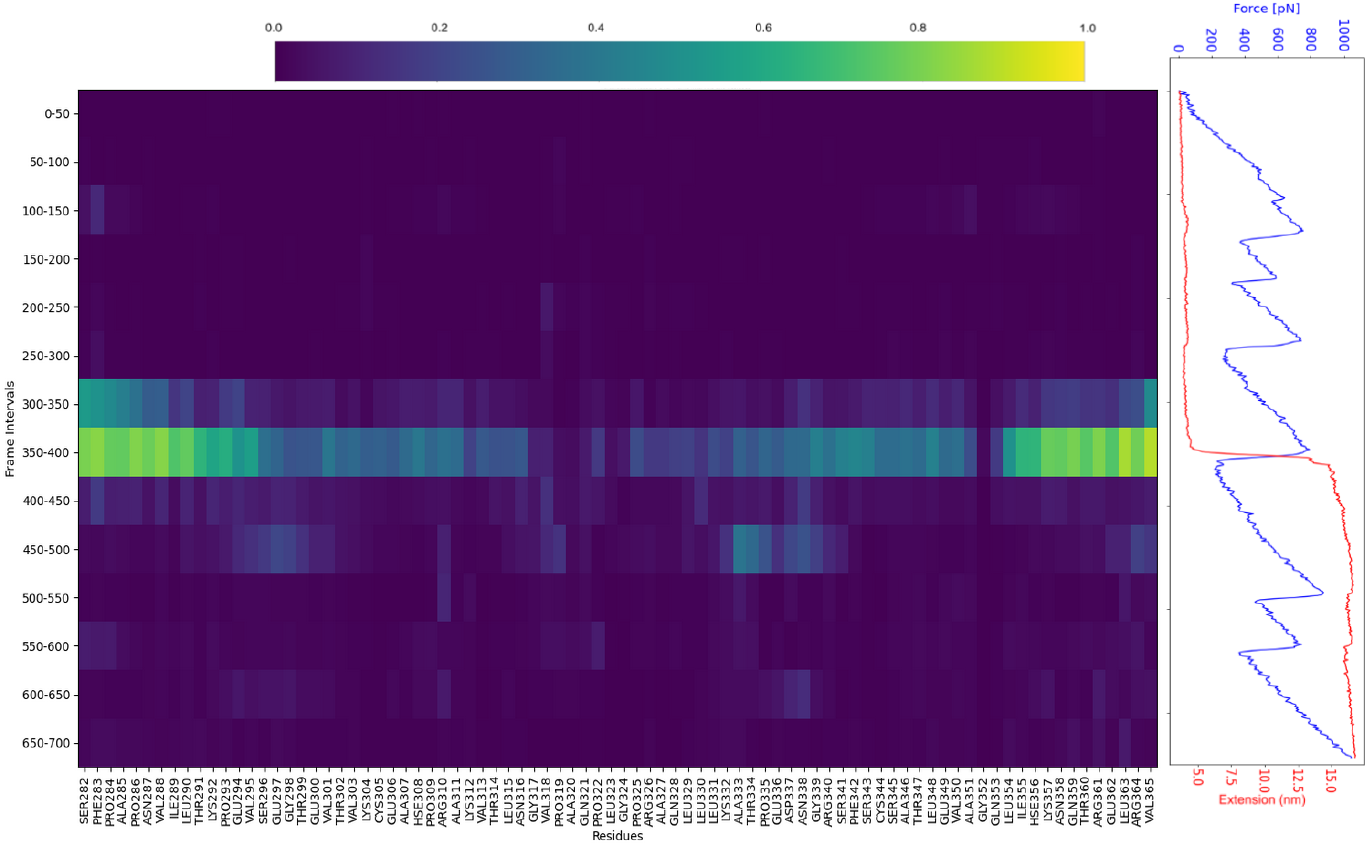
The inter-residue distance deformation measurement of the fourth domain of ICAM-1. The domain covers residues 282 to 364. The matrix is shown on the left. The calculation of the metric was performed every 50 frames. Frame intervals are reported in the *y*-axis of the matrix and the residue number is in the *x*-axis. Clusters of residues unfolding at each frame interval appear in light green and yellow. Extension of the fourth domain of ICAM-1 (red) and the total force (blue) as a function of time are shown on the right. All graphs have the same time scale.

Finally, the “see on VMD” button enables users to perform further analysis using VMD (versions 1.9 to 2.0) (10) by automatically loading the PDB and DCD files within the VMD interface installed within their local machine (**Figure S4**).

## Discussion

PySteMoDA serves as a valuable tool for the molecular dynamics community using SMD simulations with NAMD. Its primary objective is to facilitate and automate the processing and analysis of SMD simulation data. The software offers an intuitive and user-friendly interface designed to guide users at all proficiency levels, even those unfamiliar with Python. For those new to the software, a detailed tutorial is provided to ensure efficient utilization of its capabilities accessible at: https://gitlab.com/fm4b_lab/pystemoda

One notable feature of PySteMoDA is its capacity to integrate well with VMD by enabling automatic opening and data loading into it. This integration not only simplifies the workflow but also enhances the overall analytical process by providing a visual representation of the protein unfolding/unbinding trajectory.

Moreover, PySteMoDA is a semi-automatic package that offers default parameter settings that provides users with a preliminary idea of the order of magnitude for the default analysis parameters and helps understanding the characteristics of the data, without needing immediate adjustments. The software remains customizable which allows the users to modify the parameters to their specific requirements and preferences. A Google Colab (ipynb) notebook has been prepared to facilitate processing multiple simulation files, accessible via https://colab.research.google.com/drive/1Cc1IqlMnsJ-SDoE3CIiMhnLHVN5y61Dp?usp=sharing or available for download to run on a local machine as a Jupyter notebook. This Google Colab environment offers a range of advantages, including free access to a GPU runtime environment, enabling faster computations for data analysis. Furthermore, its compatibility with popular Python libraries, such as SciPy, Scikit-learn and ProDy, ensures a smooth setup for different tasks.

In general, smoothing force data facilitates the characterization of specific unfolding peaks in single-molecule SMD simulations. PySteMoDA offers two possibilities to smooth the data: natural cubic splines and Gaussian filters. In natural cubic splines, the knots parameter controls the flexibility of the spline and boundary transitions. More knots increase local control but risk overfitting by capturing noise. Properly set, knots enhance peak detection and reduce noise effectively. The Gaussian filter, on the other hand, is known for its effectiveness in signal processing (46). It attenuates noise while preserving the essential features associated with unfolding/unbinding events. Both of these smoothing approaches help users in refining peak detection.

PySteMoDA also offers two different peak detection approaches for analysis of molecular stretching pathways, the specific unfolding/unbinding peaks can be detected either from the force or from the extension curve. This second approach can be used as a complement to the first one as it may be more accurate and efficient in the identification of small peaks in some cases (e.g., in multi-domain proteins, **Figure S3**). Peak detection from the extension curve is possible via the derivative where the unfolding/unbinding peaks are reflected in sharp changes that enable a more accurate peak identification. In addition, this extension peak detection approach also allows the detection of refolding events, obscured in the force trace. These appear as steps with negative slope within the extensions versus time profile **(Figure S3A**) and can be accurately detected and exported from the derivative of the extensions profile as it is already the case with unfolding peaks. These tools accelerate the analysis of SMD simulations.

The correlation and apparent work maps enable users to identify which residues play a significant role in the mechanical stability of the protein. Maps can be calculated across a sliding window along the trajectory, providing frame specific data of pairs of residues. Residues with high correlation values within the range of frames near the unfolding/unbinding event may be critical in maintaining structural integrity under stress. Additionally, correlated residues may be evolutionarily conserved. Therefore, studying these residues (e.g., through sequence alignment with orthologs) can provide insights into the evolutionary pressure that shaped the protein’s structure and mechanical function.

The inter-residue distance deformation measurement during the trajectory is relevant to understand the dynamics of the stretched protein (**Figure 8**), as it provides insight into how the spatial relationships between residues evolve over time. For example, when the difference (*d*_*ij*_ (0) − *d*_*ij*_ (*t*)) is larger than the characteristic length scale *l*_*d*_ = 13 Å, usually used to represent 3D structures in 2D contact maps), or *l*_*d*_ = 3.8 Å, which is the typical distance between two amino acids, it indicates a significant change in the distance between residues, leading to a large exponential decay and a smaller deformation metric. This suggests that the residues have undergone a significant conformational shift (i.e., structural rearrangements). In contrast, when the distance change is smaller than *l*_*d*_, the exponential term will be closer to 1, and the deformation metric will be larger, indicating minimal change. The subtraction from 1 at the start makes the metric more intuitive. Therefore, the higher the deformation the closer the metric is to 1. This method has also been adapted to compute deformation in terms of work (in Joules), **see supp. information**. These calculations can be performed using the provided Google Colab notebook or via the command line, while the analysis of these spatial relationships gives insight into the conformational rearrangement of the molecule before and after unfolding events.

## Conclusion

PySteMoDA finds application in computational protein dynamics where researchers are interested in protein unfolding and unbinding. Researchers can use PySteMoDA to extract and analyze forces, trajectories, and structural data from SMD simulations which enables them to gain deeper insights into the predicted behavior of biomolecules. PySteMoDA represents a valuable addition to the toolkit of computational biology and researchers involved in using SMD simulations. Its user-friendly Graphical User Interface (GUI), data handling capabilities, open-source nature and comprehensive documentation make it accessible to new and experienced users.

## Data availability

The package is free for all users: https://pypi.org/project/PySteMoDA/0.1.0/

The code is hosted on GitLab:https://gitlab.com/fm4b_lab/pystemoda

Users who do not have python3 within their local machine can use the Google Colab to avoid the installation process: https://colab.research.google.com/drive/1Cc1IqlMnsJ-SDoE3CIiMhnLHVN5y61Dp?usp=sharing

## Acknowledgement

The authors acknowledge Dr. Jean-Michel Arbona for insightful discussions on the inter-residue deformation measurement. Some simulations were carried out at the Ohio Supercomputing Center.

## References

1. Smith JC, Roux B. Eppur si muove! The 2013 Nobel Prize in Chemistry. Struct Lond Engl 1993. 2013 Dec 3;21(12):2102–5.

2. Dror RO, Dirks RM, Grossman JP, Xu H, Shaw DE. Biomolecular simulation: a computational microscope for molecular biology. Annu Rev Biophys. 2012;41:429–52.

3. Lee EH, Hsin J, Sotomayor M, Comellas G, Schulten K. Discovery through the computational microscope. Struct Lond Engl 1993. 2009 Oct 14;17(10):1295–306.

4. Adcock SA, McCammon JA. Molecular dynamics: survey of methods for simulating the activity of proteins. Chem Rev. 2006 May;106(5):1589–615.

5. Karplus M, Petsko GA. Molecular dynamics simulations in biology. Nature. 1990 Oct;347(6294):631–9.

6. Daggett V, Levitt M. Realistic simulations of native-protein dynamics in solution and beyond. Annu Rev Biophys Biomol Struct. 1993;22:353–80.

7. Case DA, Cheatham III TE, Darden T, Gohlke H, Luo R, Merz Jr. KM, et al. The Amber biomolecular simulation programs. J Comput Chem. 2005;26(16):1668–88.

8. Abraham MJ, Murtola T, Schulz R, Páll S, Smith JC, Hess B, et al. GROMACS: High performance molecular simulations through multi-level parallelism from laptops to supercomputers. SoftwareX. 2015 Sep 1;1–2:19–25.

9. Phillips JC, Braun R, Wang W, Gumbart J, Tajkhorshid E, Villa E, et al. Scalable Molecular Dynamics with NAMD. J Comput Chem. 2005 Dec;26(16):1781–802.

10. Humphrey W, Dalke A, Schulten K. VMD: visual molecular dynamics. J Mol Graph. 1996 Feb;14(1):33–8, 27–8.

11. Isralewitz B, Gao M, Schulten K. Steered molecular dynamics and mechanical functions of proteins. Curr Opin Struct Biol. 2001 Apr 1;11(2):224–30.

12. Franz F, Daday C, Gräter F. Advances in molecular simulations of protein mechanical properties and function. Curr Opin Struct Biol. 2020 Apr 1;61:132–8.

13. Sotomayor M, Schulten K. Single-molecule experiments in vitro and in silico. Science. 2007 May 25;316(5828):1144–8.

14. Grubmüller H. Force probe molecular dynamics simulations. Methods Mol Biol Clifton NJ. 2005;305:493–515.

15. Grubmüller H, Heymann B, Tavan P. Ligand binding: molecular mechanics calculation of the streptavidin-biotin rupture force. Science. 1996 Feb 16;271(5251):997–9.

16. Lu H, Isralewitz B, Krammer A, Vogel V, Schulten K. Unfolding of Titin Immunoglobulin Domains by Steered Molecular Dynamics Simulation. Biophys J. 1998 Aug 1;75(2):662–71.

17. Park S, Schulten K. Calculating potentials of mean force from steered molecular dynamics simulations. J Chem Phys. 2004 Apr 1;120(13):5946–61.

18. Izrailev S, Stepaniants S, Balsera M, Oono Y, Schulten K. Molecular dynamics study of unbinding of the avidin-biotin complex. Biophys J. 1997 Apr;72(4):1568–81.

19. Rico F, Russek A, González L, Grubmüller H, Scheuring S. Heterogeneous and rate-dependent streptavidin–biotin unbinding revealed by high-speed force spectroscopy and atomistic simulations. Proc Natl Acad Sci. 2019 Apr 2;116(14):6594–601.

20. Milles LF, Gaub HE. Extreme mechanical stability in protein complexes. Curr Opin Struct Biol. 2020 Feb;60:124–30.

21. Lu H, Schulten K. The Key Event in Force-Induced Unfolding of Titin’s Immunoglobulin Domains. Biophys J. 2000 Jul 1;79(1):51–65.

22. Evans E, Ritchie K. Dynamic strength of molecular adhesion bonds. Biophys J. 1997 Apr;72(4):1541–55.

23. Marszalek PE, Lu H, Li H, Carrion-Vazquez M, Oberhauser AF, Schulten K, et al. Mechanical unfolding intermediates in titin modules. Nature. 1999 Nov 4;402(6757):100–3.

24. Rico F, Gonzalez L, Casuso I, Puig-Vidal M, Scheuring S. High-Speed Force Spectroscopy Unfolds Titin at the Velocity of Molecular Dynamics Simulations. Science. 2013 Nov 8;342(6159):741–3.

25. Nisler CR, Narui Y, Scheib E, Choudhary D, Bowman JD, Mandayam Bharathi H, et al. Interpreting the Evolutionary Echoes of a Protein Complex Essential for Inner-Ear Mechanosensation. Mol Biol Evol. 2023 Apr 4;40(4):msad057.

26. Upadhyaya A, Baraban M, Wong J, Matsudaira P, van Oudenaarden A, Mahadevan L. Power-limited contraction dynamics of Vorticella convallaria: an ultrafast biological spring. Biophys J. 2008 Jan 1;94(1):265–72.

27. Jaiganesh A, Narui Y, Araya-Secchi R, Sotomayor M. Beyond Cell-Cell Adhesion: Sensational Cadherins for Hearing and Balance. Cold Spring Harb Perspect Biol. 2018 Sep 4;10(9):a029280.

28. Robles L, Ruggero MA. Mechanics of the mammalian cochlea. Physiol Rev. 2001 Jul;81(3):1305–52.

29. Corey DP, Hudspeth AJ. Kinetics of the receptor current in bullfrog saccular hair cells. J Neurosci Off J Soc Neurosci. 1983 May;3(5):962–76.

30. Freddolino PL, Liu F, Gruebele M, Schulten K. Ten-microsecond molecular dynamics simulation of a fast-folding WW domain. Biophys J. 2008 May 15;94(10):L75–77.

31. Sotomayor M. Computational exploration of single-protein mechanics by steered molecular dynamics. AIP Conf Proc. 2015 Dec 31;1703(1):030001.

32. Costescu BI, Sturm S, Gräter F. Dynamic disorder can explain non-exponential kinetics of fast protein mechanical unfolding. J Struct Biol. 2017 Jan 1;197(1):43–9.

33. Orzeł U, Pasznik P, Miszta P, Lorkowski M, Niewieczerzał S, Jakowiecki J, et al. GS-SMD server for steered molecular dynamics of peptide substrates in the active site of the γ-secretase complex. Nucleic Acids Res. 2023 Jul 5;51(W1):W251–62.

34. Bernardi RC, Durner E, Schoeler C, Malinowska KH, Carvalho BG, Bayer EA, et al. Mechanisms of Nanonewton Mechanostability in a Protein Complex Revealed by Molecular Dynamics Simulations and Single-Molecule Force Spectroscopy. J Am Chem Soc. 2019 Sep 18;141(37):14752–63.

35. Bezerra Beniz D, Espíndola A. USING TKINTER OF PYTHON TO CREATE GRAPHICAL USER INTERFACE (GUI) FOR SCRIPTS IN LNLS. 2016.

36. Bakan A, Meireles LM, Bahar I. ProDy: Protein Dynamics Inferred from Theory and Experiments. Bioinformatics. 2011 Jun 1;27(11):1575–7.

37. Pedregosa F, Varoquaux G, Gramfort A, Michel V, Thirion B, Grisel O, et al. Scikit-learn: Machine Learning in Python. J Mach Learn Res. 2011 Nov 1;12(null):2825–30.

38. Virtanen P, Gommers R, Oliphant TE, Haberland M, Reddy T, Cournapeau D, et al. SciPy 1.0: fundamental algorithms for scientific computing in Python. Nat Methods. 2020 Mar;17(3):261–72.

39. Jaiganesh A, De-la-Torre P, Patel AA, Termine DJ, Velez-Cortes F, Chen C, et al. Zooming in on cadherin-23: Structural diversity and potential mechanisms of inherited deafness. Struct Lond Engl 1993. 2018 Sep 4;26(9):1210-1225.e4.

40. Sotomayor M, Weihofen WA, Gaudet R, Corey DP. Structural Determinants of Cadherin-23 Function in Hearing and Deafness. Neuron. 2010 Apr 15;66(1):85–100.

41. Vanommeslaeghe K, Hatcher E, Acharya C, Kundu S, Zhong S, Shim J, et al. CHARMM General Force Field (CGenFF): A force field for drug-like molecules compatible with the CHARMM all-atom additive biological force fields. J Comput Chem. 2010 Mar;31(4):671–90.

42. Hwang W, Austin SL, Blondel A, Boittier ED, Boresch S, Buck M, et al. CHARMM at 45: Enhancements in Accessibility, Functionality, and Speed. J Phys Chem B. 2024 Oct 17;128(41):9976–10042.

43. Xiao C, Bator-Kelly CM, Rieder E, Chipman PR, Craig A, Kuhn RJ, et al. The crystal structure of coxsackievirus A21 and its interaction with ICAM-1. Struct Lond Engl 1993. 2005 Jul;13(7):1019–33.

44. Chen X, Kim TD, Carman CV, Mi LZ, Song G, Springer TA. Structural plasticity in Ig superfamily domain 4 of ICAM-1 mediates cell surface dimerization. Proc Natl Acad Sci. 2007 Sep 25;104(39):15358–63.

45. Feng G. Data smoothing by cubic spline filters. Signal Process IEEE Trans On. 1998 Nov 1;46:2790–6.

46. Deisenroth M, Ohlsson H. A general perspective on Gaussian filtering and smoothing: Explaining current and deriving new algorithms. 2011. 1807 p.

