## Supplementary material for "PySteMoDA: An Open-Source Python Package for the Analysis of Steered Molecular Dynamics Simulations Data": supp_material

### Data extraction and conversion

To obtain the SMD forces in the direction of pulling in picoNewton (pN), PySteMoDA extracts raw SMD forces (kcal/mol/Å) and the direction of pulling. Then the dot product of both of these parameters is calculated:

$$F(v) \ = \ F_{smd}(Fx,\ Fy,\ Fz) \cdot direction_{pulling}(x,\ y,\ z)$$

where the resulting force is in pN.

If any TCL forces (kcal/mol/Å) have been applied during the simulation and recorded in the LOG file, PySteMoDA extracts them automatically and converts them to picoNewton:

$$F(-\ \nu) \ = \ F_{tcl} \times 69.479$$

assuming 1 kcal/mol = 69.479 pN· Å

The spring constant recorded in kcal/mol/Å² is converted to pN/nm:

$$k_{(pN/nm)} \ = \ k_{(kcal/mol/Å^2)} \times 69.479 \times 10$$

The time is converted to seconds knowing the time step of the simulation, in most of the cases corresponding to 2 femtoseconds, in our simulations:

$$Time_{(seconds)} \ = \ TimeStep \times 2 \times 10^{-15}$$

| Peak | timeStep | timeInS | UnfoldingForce [pN] | UnfoldingLoadingRate [pN/s] | Stiffness [pN/nm] | UnfoldingDisplacement [nm] | InterDomain Distance [nm] |
|---|---|---|---|---|---|---|---|
| 1 | 547,500 | 1.095e-09 | 1,986 | 2507649530253.981 | 501.52 | 5.5 | 5.2 |
| 2 | 1,067,500 | 2.135e-09 | 1,045 | 1849659473948.7783 | 369.93 | 10.7 | |

**Table S1:** Data export of the peak detection results shown in Figure 4. Users have the possibility to select the peaks of their interest and export them to files of different formats (csv, txt, dat).

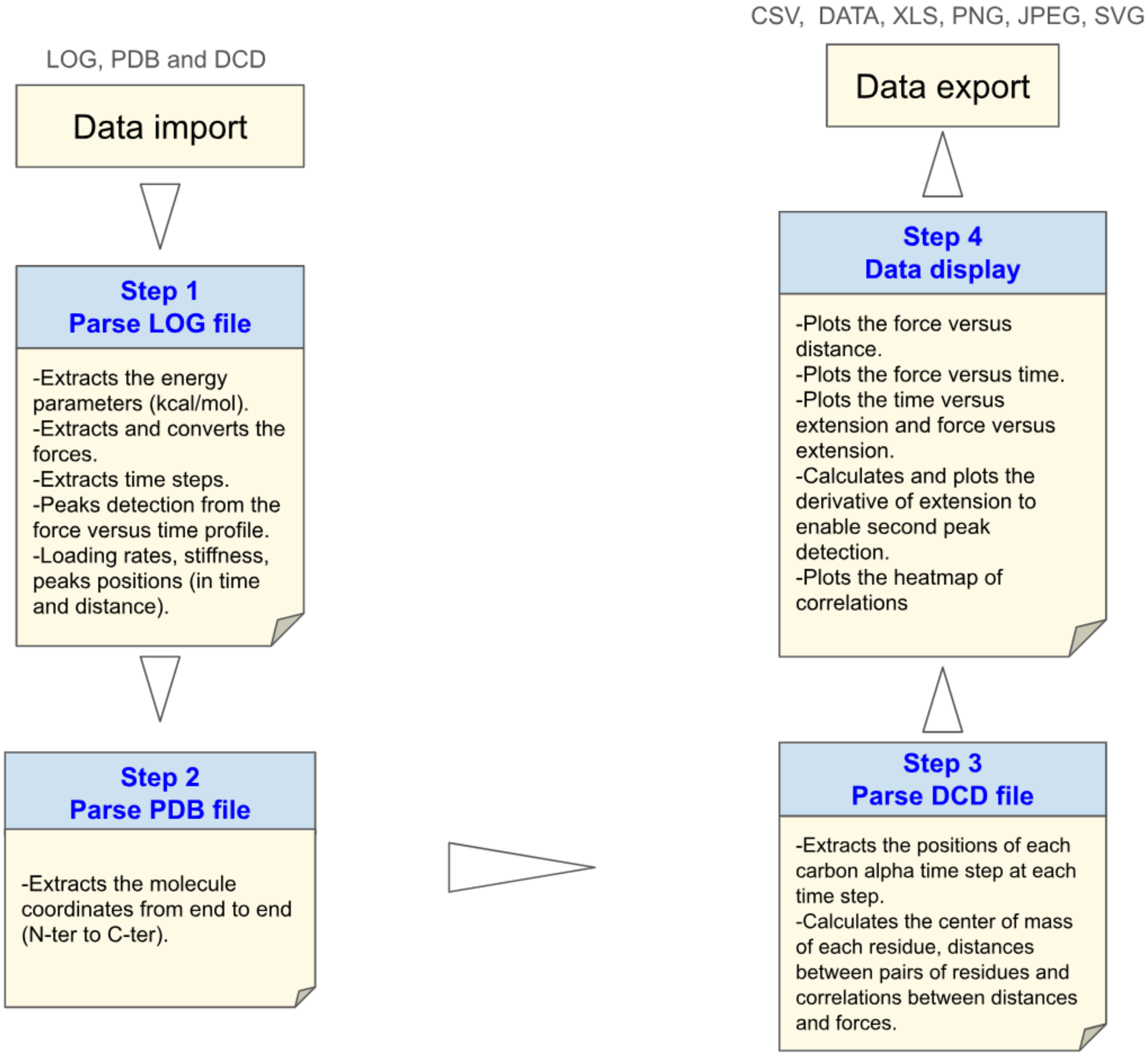


**Figure S1:** A schematic review of the PySteMoDA pipeline. Through each step PySteMoDA extracts and converts (if needed) the data. Default parameters are automatically suggested and used (e.g; smoothing and peak detection). These parameters can be modified by the user. At the end of the analysis, users have the possibility to export their data and plots in different formats.

A

B

C

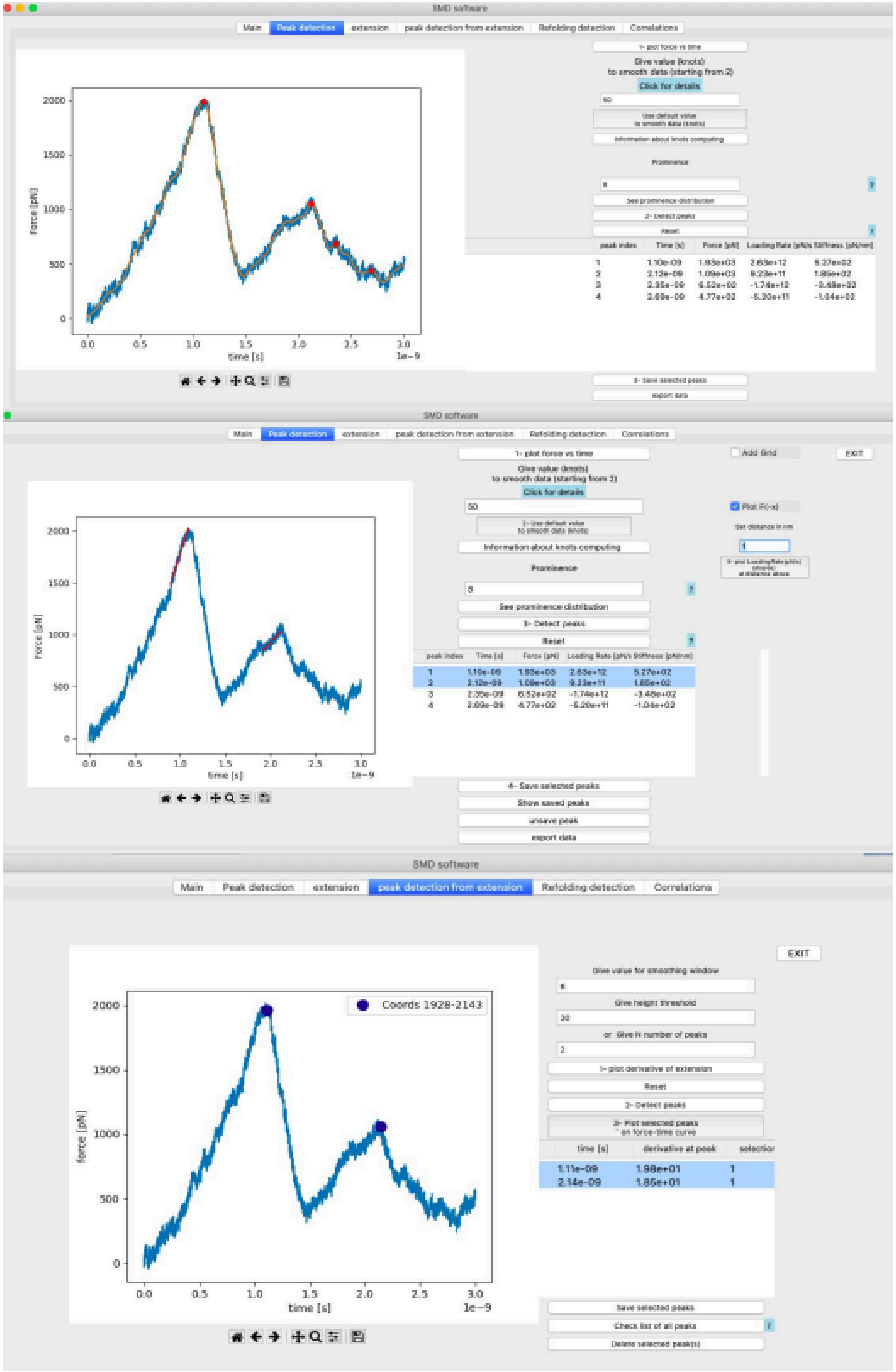

**Figure S2:** Snapshot of the TCL forces curve for CDH23 EC19-20. **(A)** data smoothing and peak detection. **(B-C)** Loading rates at each force peak and (B) and peak detection from the extension curve via the derivative (C).

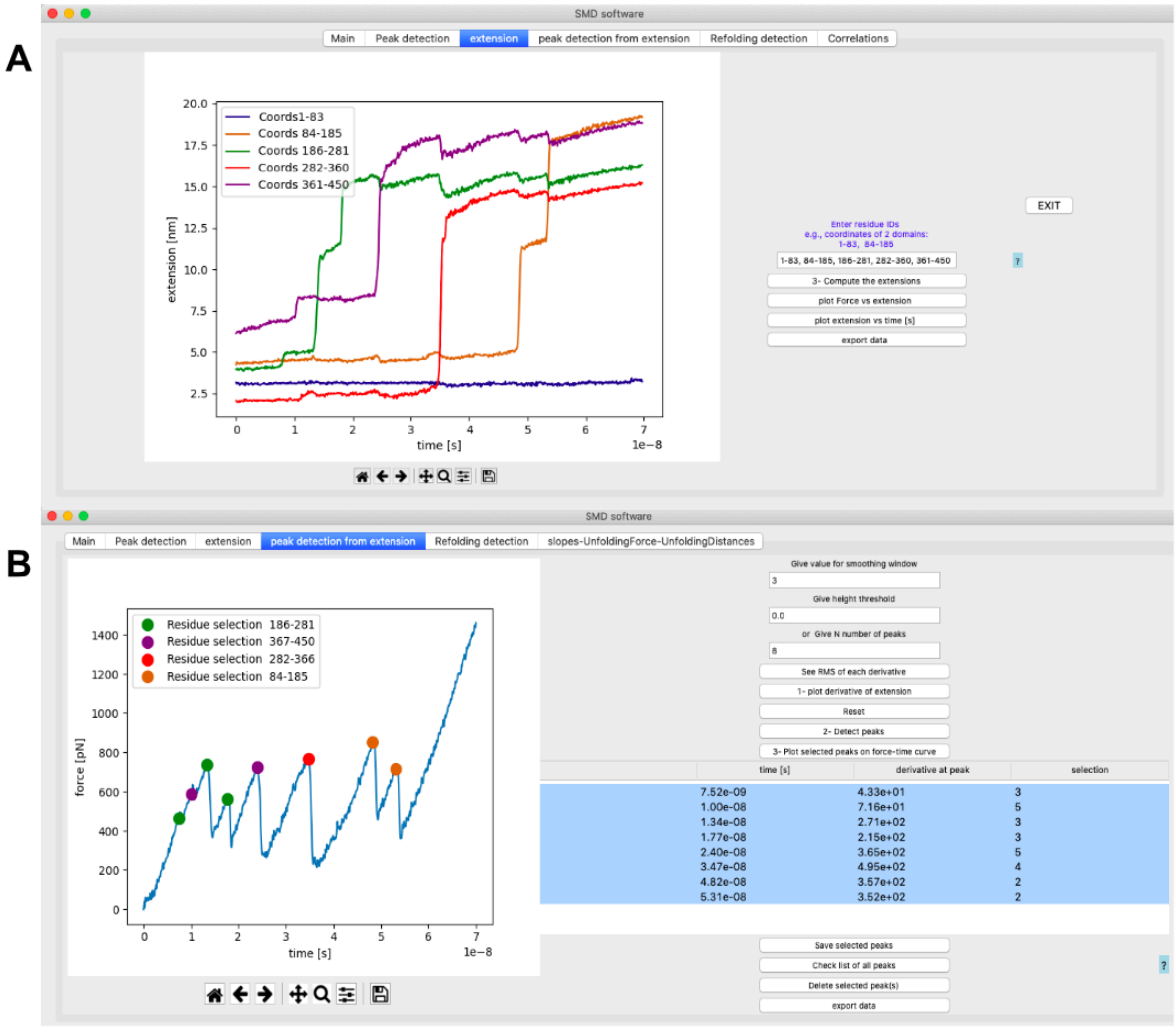


**Figure S3:** PySteMoDA analysis of the SMD trajectory for five domains of the human intercellular adhesion molecule 1 (ICAM-1). **(A)** extensions of each domain with respect to time; the specific unfolding events appear as increasing steps whereas the refolding events appear as small decreasing steps (see green and purple curves). **(B)** peak detection from the derivative of the extension curve, this approach allows users to detect which domain unfolded at which force peak. ICAM-1 simulations were carried out in the *NVE* ensemble at 1 nm/ns.

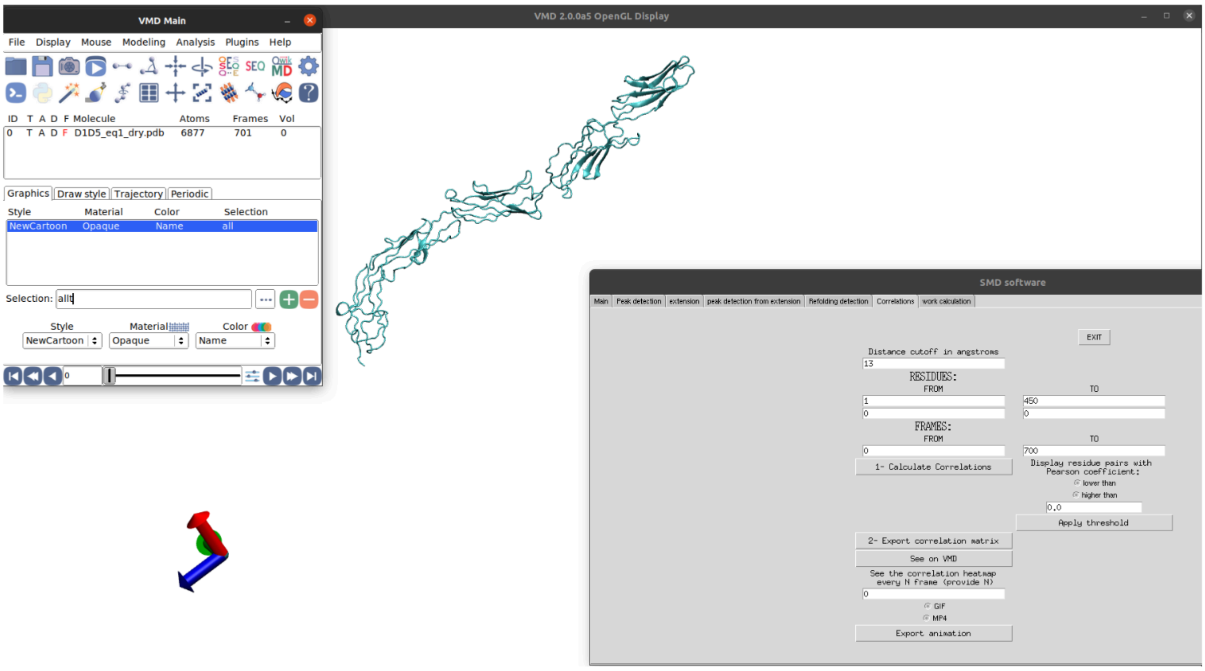


**Figure S4:** A snapshot of PySTeMoDA's graphical user interface (GUI) showing its integration with VMD. By simply pressing the "See on VMD" button, the software launches a VMD window showing the protein trajectory during unfolding.
